# Adrenomedullin-CALCRL Axis Controls Relapse-Initiating Drug Tolerant Acute Myeloid Leukemia Cells

**DOI:** 10.1101/2020.08.17.253542

**Authors:** Clément Larrue, Nathan Guiraud, Pierre-Luc Mouchel, Marine Dubois, Thomas Farge, Mathilde Gotanègre, Claudie Bosc, Estelle Saland, Marie-Laure Nicolau-Travers, Marie Sabatier, Nizar Serhan, Ambrine Sahal, Emeline Boet, Sarah Mouche, Quentin Heydt, Nesrine Aroua, Lucille Stuani, Tony Kaoma, Linus Angenendt, Jan-Henrik Mikesch, Christoph Schliemann, François Vergez, Jérôme Tamburini, Christian Récher, Jean-Emmanuel Sarry

## Abstract

Drug tolerant leukemic cell subpopulations may explain frequent relapses in acute myeloid leukemia (AML), suggesting that these Relapse-Initiating Cells (RICs) persistent after chemotherapy represent *bona fide* targets to prevent drug resistance and relapse. We uncovered that the G-protein coupled receptor CALCRL is expressed in leukemic stem cells (LSCs) and RICs, and that the overexpression of CALCRL and/or of its ligand adrenomedullin (ADM) and not CGRP correlates to adverse outcome in AML. CALCRL knockdown impairs leukemic growth, decreases LSC frequency and sensitizes to cytarabine in patient-derived xenograft (PDX) models. Mechanistically, the ADM-CALCRL axis drives cell cycle, DNA repair and mitochondrial OxPHOS function of AML blasts dependent on E2F1 and BCL2. Finally, CALCRL depletion reduces LSC frequency of RICs post-chemotherapy *in vivo*. In summary, our data highlight a critical role of ADM-CALCRL in post-chemotherapy persistence of these cells, and disclose a promising therapeutic target to prevent relapse in AML.

---

Despite improvements in the complete response rate obtained after conventional chemotherapy, the overall survival of acute myeloid leukemia (AML) patients is still poor, due to frequent relapses caused by chemotherapy resistance. Although novel targeted therapies are holding great promises, eradicating drug tolerant/resistant AML cells after chemotherapy remains the major challenge in the treatment of AML. AML arises from self-renewing leukemic stem cells (LSCs) that can repopulate human AML when assayed and xenografted in immunocompromised mice (e.g. PDX; Bonnet and Dick, 1997). Even though these cells account for a minority of leukemic burden, gene signatures associated with a stem cell phenotype or function have an unfavorable prognosis in AML (Eppert et al., 2011; Gentles, 2010; Ng et al., 2016; Vergez et al., 2011; Zeijlemaker et al., 2018). Clinical relevance is evidenced by the enrichment in Leukemia-Initiating Cells (LICs) and LSC-related gene signatures in AML specimens at the time of relapse compared to diagnosis (Hackl et al., 2015; Ho et al., 2016). While it has been initially shown that LSCs may be less affected by chemotherapy than more mature populations (Jordan et al., 2006; Ishikawa et al., 2007; Thomas et al. 2013), recent works demonstrated that the anti-AML chemotherapy cytarabine (AraC) might deplete the LSC pool in PDX models (Farge et al., 2017; Boyd et al., 2018). These results suggest the coexistence of two distinct LSC populations, one chemosensitive and thus eradicated by conventional treatments, and one that is chemoresistant, persists and might induce relapse in AML (RIC for Relapse-initiating drug tolerant Cells). Thus, a better phenotypical and functional characterization of RICs is crucial for the development of new AML therapies aiming at reducing the risk of relapse.

Although it was first proposed that the LSC-compartment was restricted to the CD34^+^CD38^-^ subpopulation of human AML cells (Bonnet and Dick, 1997; Ishikawa et al., 2007), several studies subsequently demonstrated that LSCs are phenotypically heterogeneous when assayed in NSG mice (Taussig et al., 2010; Sarry et al., 2011; Eppert et al., 2011; Quek et al., 2016). These observations highlight that more functional studies are needed to better and more fully characterize LSCs particularly under the selection pressure imposed by chemotherapy. Eradicating LSCs without killing normal hematopoietic stem cells (HSCs) depends on the identification of therapeutically relevant markers that are overexpressed in the AML compartment. In recent years, tremendous research efforts led to the identification of several cell surface markers such as CD47, CD123, CD44, TIM-3, CD25, CD32 or CD93 discriminating LSCs from HSCs (Iwasaki et al., 2015; Jin et al., 2009; Kikushige et al., 2010; Majeti et al., 2009; Perna et al., 2017; Saito et al., 2010). In addition, it has been proposed that LSCs also have both a specific decrease in their reactive oxygen species (ROS) content and an increase in BCL2-dependent oxidative phosphorylation (OxPHOS), revealing a vulnerability that can be exploited through treatment with BCL2 inhibitors such as venetoclax (Lagadinou et al., 2013; Konopleva et al., 2016). This is consistent with several studies demonstrating that mitochondrial OxPHOS status contributes to drug resistance in myeloid leukemia (Bosc et al., 2017; Farge et al., 2017; Henkenius et al., 2017; Kuntz et al., 2017). Taken together, these results suggest that specific characteristics of LSCs can be exploited to develop targeted therapeutic approaches.

Here, we report that the cell surface G protein-coupled receptor (GPCR) family calcitonin receptor-like receptor (CALCRL) and its ligand adrenomedullin (ADM) are expressed in AML cells and that CALCRL sustains LSC function. High expression of both CALCRL and ADM is predictive of an unfavorable prognosis in a cohort of 179 AML patients. Moreover, depletion of CALCRL abrogates leukemic growth and dramatically induces cell death *in vivo*. Transcriptomic and functional analyses show that CALCRL drives E2F1, BCL2 and OxPHOS pathways involved in the chemoresistance. Furthermore, we observed that cell surface expression of CALCRL is enriched after AraC treatment in PDX models as well as after intensive chemotherapy in AML patients. Limiting dilution analyses coupled to genetic manipulation demonstrate that RICs are critically dependent on CALCRL for their maintenance. Altogether, our findings demonstrate that CALCRL is a new RIC player with a critical effector role in both their stemness and chemoresistance, and thus is a relevant target to eradicate this specific cell population.

## Results

### The receptor CALCRL and its ligand adrenomedullin are expressed in AML cells and associated with a poor outcome in patients

Using a clinically relevant chemotherapeutic model, we and others previously demonstrated that LSCs are not necessarily enriched in post-AraC residual AML, suggesting LSCs include both chemosensitive and chemoresistant stem cell sub-populations (Farge et al., 2017; Boyd et al., 2018). In order to identify new vulnerabilities in the chemoresistant LSC population that might be responsible for relapse, we analyzed transcriptomic data from three different studies that (Figure 1A; Table S1): i) identified 134 genes overexpressed in functionally defined LSC compared with a normal HSC counterpart (Eppert et al., 2011; GSE30377); ii) uncovered 114 genes of high expression associated with poor prognosis in AML (the Cancer Genome Atlas, AML cohort, 2013); and iii) selected 536 genes overexpressed at relapse compared to pairwise matched diagnosis samples (Hackl et al., 2015; GSE66525). Surprisingly, we found one unique gene common to these three independent transcriptomic datasets: *CALCRL*, encoding a G protein-coupled seven-transmembrane domain receptor poorly documented in cancer that has been recently described as associated with a poor prognosis in AML (Angenendt et al, 2019). Using four independently published cohorts of AML patients (TCGA AML cohort; GSE12417; GSE14468; BeatAML cohort), we observed that patients with high *CALCRL* expression had a shorter overall survival (Figure 1B; Figure S1A) and are more refractory to chemotherapy (Figure S1B) compared to patients with low *CALCRL* expression. This correlated with a higher expression in complex *versus* normal karyotypes (Figure S1C). Furthermore, *CALCRL* gene expression was significantly higher at relapse compared to diagnosis in patients treated with intensive chemotherapy (Figure 1C). *CALCRL* expression was also higher in the leukemic compartment compared with normal hematopoietic cells, and more specifically in the LSC population as both functionally-(Figure 1D) and phenotypically- (Figure S1D) defined compared with the AML bulk population. Interestingly, *CALCRL* expression was higher in the more immature AML subtypes according to FAB stratification, suggesting that CALCRL is a marker of cell immaturity (Figure S1E). Using flow cytometry, we determined that CALCRL was expressed at the cell surface (Figure S1F), more markedly in leukemic compared to normal CD34+ hematopoietic progenitor cells (Figure S1G; see Table S2 for mutational status of patients). Of note, *CALCRL* expression did not correlate with any most found mutations (Figure S1). Next, we assessed the expression of ADM, a CALCRL ligand already described in several solid cancers (Berenguer-Daizé et al., 2013; Kocemba et al., 2013). The *ADM* gene is overexpressed in AML cells compared to normal cells (Figure S1J-K) although its expression is not altered in AML patients at relapse compared to diagnosis (Figure S1L) and is not linked to mutational status (Figure S1I). Using a combination of western blotting, confocal microscopy and RNA microarray, we have established that CALCRL, its three co-receptors RAMP1, RAMP2 and RAMP3, as well as ADM (but not CGRP, another putative CALCRL ligand) are expressed in all tested AML cell lines and primary AML samples (Figure S1M-R). Moreover, analysis of three independent cohorts (TCGA, Verhaak *et al*. and BeatAML) confirmed that only *ADM* was highly expressed in primary samples compared to *CALCA* and *CALCB* (two genes encoding CGRP) that were not or poorly expressed (Figure S1S). Next, we addressed the impact of CALCRL and ADM protein levels at diagnosis on patient outcome. Using IHC analyses, we observed that increasing protein levels of CALCRL or ADM were associated with decreasing complete remission rates, inferior 5-year overall and event-free survival in a cohort of 179 intensively treated AML patients (Figure 1E and 1F). When patients were clustered into 4 groups according to CALCRL and ADM expression (low/low *vs* low/high *vs* high/low *vs* high/high; Figure 1G; Table S3), we observed that the CALCRL^high^/ADM^high^ group was associated with the lowest overall survival and that high expression of only CALCRL or ADM also correlated to reduced EFS and complete remission rate (Figure 1H). Next, we addressed the protein level of CGRP using IHC analyses and we detected a slight diffuse signal of this protein, suggesting a paracrine secretion of CGRP in AML. However, we demonstrated that protein levels of CGRP had no impact on 5-year overall and event-free survival (Figure S2A-B-C), indicating that ADM was likely the main driver of CALCRL activation in AML.

**Figure 1.**
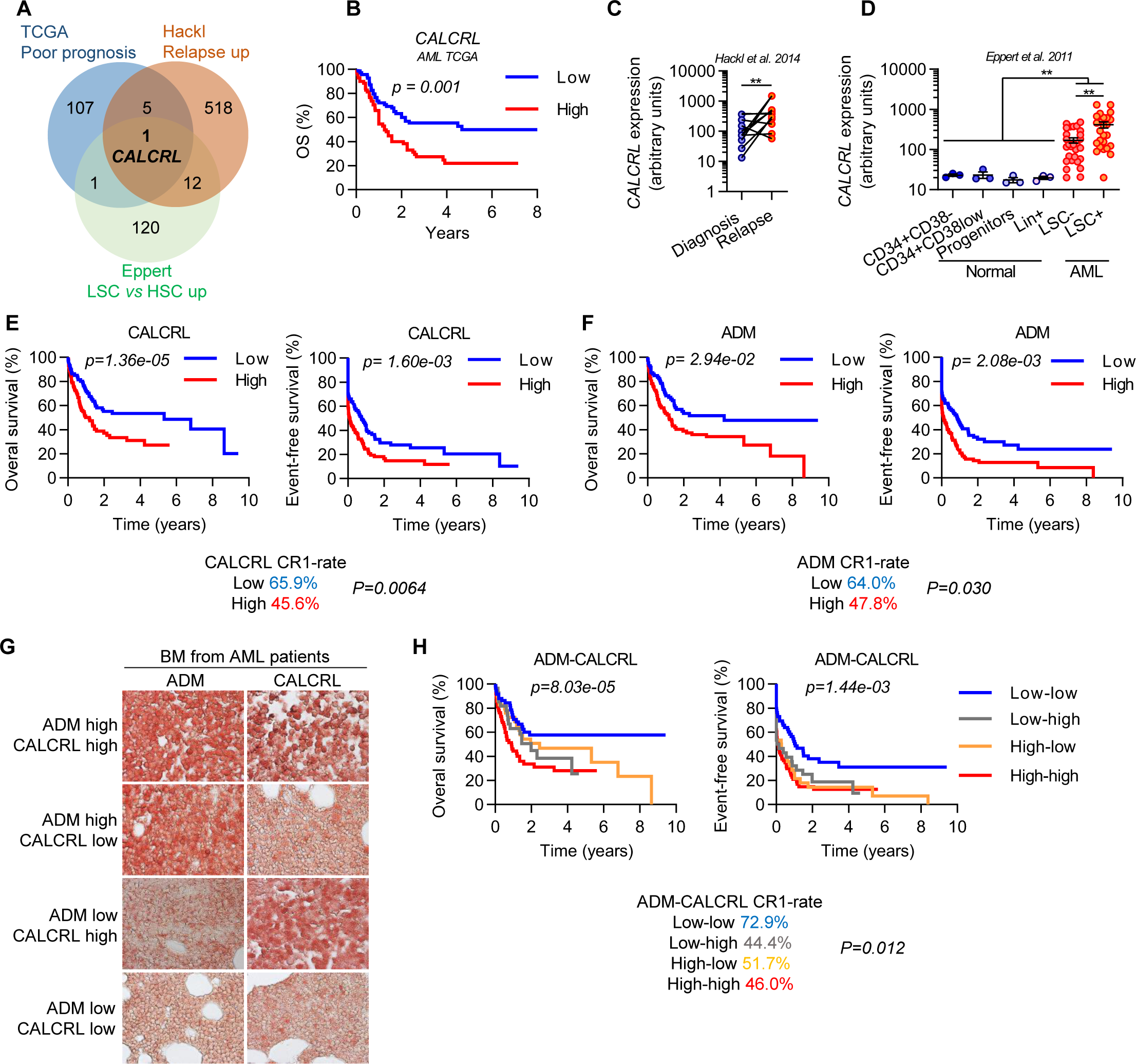
Expression of CALCRL and its ligand Adrenomedullin and impact on patient outcome in AML. (A) Genes overexpressed and associated with poor prognosis, relapse or LSC *vs* HSC according to TCGA, Hackl *et al*. and Eppert *et al*. studies respectively. (B) Impact of *CALCRL* gene expression on overall survival in the TCGA cohort. Data were dichotomized according to median value for visualization purpose. (C) *CALCRL* expression at diagnosis and relapse after intensive chemotherapy according to Hackl *et al*. (D) *CALCRL* gene expression in normal compartments CD34^+^CD38^-^, CD34^+^CD38^low^, progenitors and Lin^+^ compared with leukemic LSC^-^ and LSC^+^ compartments according to Eppert *et al*. (E-F) Overall survival (OS) and Event-Free Survival (EFS) according to CALCRL and ADM H-scores. OS and EFS were regressed against CALCRL and/or ADM H-scores using univariate or multivariate Cox model. Data were dichotomized for visualization purpose. (G) Representative IHC micrographs of CALCRL and ADM expression in pretherapeutic BM from AML patients. (H) Overall and Event-Free Survival according to CALCRL and ADM H-scores. OS and EFS were regressed against CALCRL and/or ADM H-scores using univariate or multivariate Cox model. Data were dichotomized for visualization purpose. *p<0.05; **p<0.01; ***p<0.001; ns, not significant.

All these data supported the hypothesis that the ADM-CALCRL axis is activated in an autocrine dependent manner and associated with a poor prognosis in AML.

### The CALCRL-ADM axis is required for cell growth and survival

Next, we investigated the impact of deregulated CALCRL-ADM axis in cell proliferation and survival. CALCRL depletion was associated with a decrease of blast cell proliferation (Figure 2A), and an increase in cell death (Figure 2B) in three AML (MOLM-14, OCI-AML2, OCI-AML3) cell lines. Furthermore, ADM-targeting shRNA (Figure S3A) phenocopied the effects of shCALCRL on cell proliferation and apoptosis in MOLM-14 and OCI-AML3 cells (Figure S3B-C). In order to confirm these results *in vivo* and to control the invalidation of the target over time, we have developed tetracycline-inducible shRNA models. First, we established that the inducible depletion of CALCRL was associated with a decrease in cell proliferation and an increase in apoptosis as observed with constitutive shRNA approaches (Figure S3D-E-F). After injection of AML cells in mice, RNA depletion was activated from the first day by doxycycline (Figure 2C). Twenty-five days post-transplantation, the engraftment of human leukemic cells from murine bone marrow and spleen was assessed with mCD45.1^-^hCD45^+^hCD33^+^AnV^-^ markers (Figure S3G). Mice injected with shCAL#1 and shCAL#2 had a significant reduction in total cell tumor burden (as defined by AML blasts in the bone marrow + spleen) compared to shCTR (CTR for control) in both MOLM-14 (shCTR=13.9×10^6^ cells *vs* shCAL#1=0.3x10^6^ cells *vs* shCAL#2=0.1×10^6^ cells) and OCI-AML3 cells (shCTR=17.2×10^6^ cells *vs* shCAL#1=2.0×10^6^ cells *vs* shCAL#2=1.7×10^6^ cells) (Figure 2D; Figure S3H). Finally, CALCRL silencing significantly prolonged mice survival (Figure 2E). To take full advantage of our inducible constructs and to improve the clinical relevance of our model, we assessed the impact of CALCRL depletion on highly engrafting AML cells (Figure 2F). Short hairpin RNA expression was induced 10 days post-transplantation of shCTR or shCAL in MOLM14 cells after verifying that the level of engraftment was similar in both groups (Figure 2F-G). In this overt AML model, we observed a marked reduction in bone marrow blasts in the mice xenografted with shCAL AML cells compared to the shCTR-xenografted cohort in which the disease progressed (Figure 2G). Furthermore, CALCRL downregulation significantly increased survival (Figure 2H). Importantly, these results demonstrated that the reduction of blast number and the increased survival of mice observed after CALCRL depletion was not the consequence of an inhibition of leukemic blast homing to the murine bone marrow. Altogether, these results demonstrate that CALCRL is required both for the propagation and the maintenance of AML cells *in vivo*.

**Figure 2.**
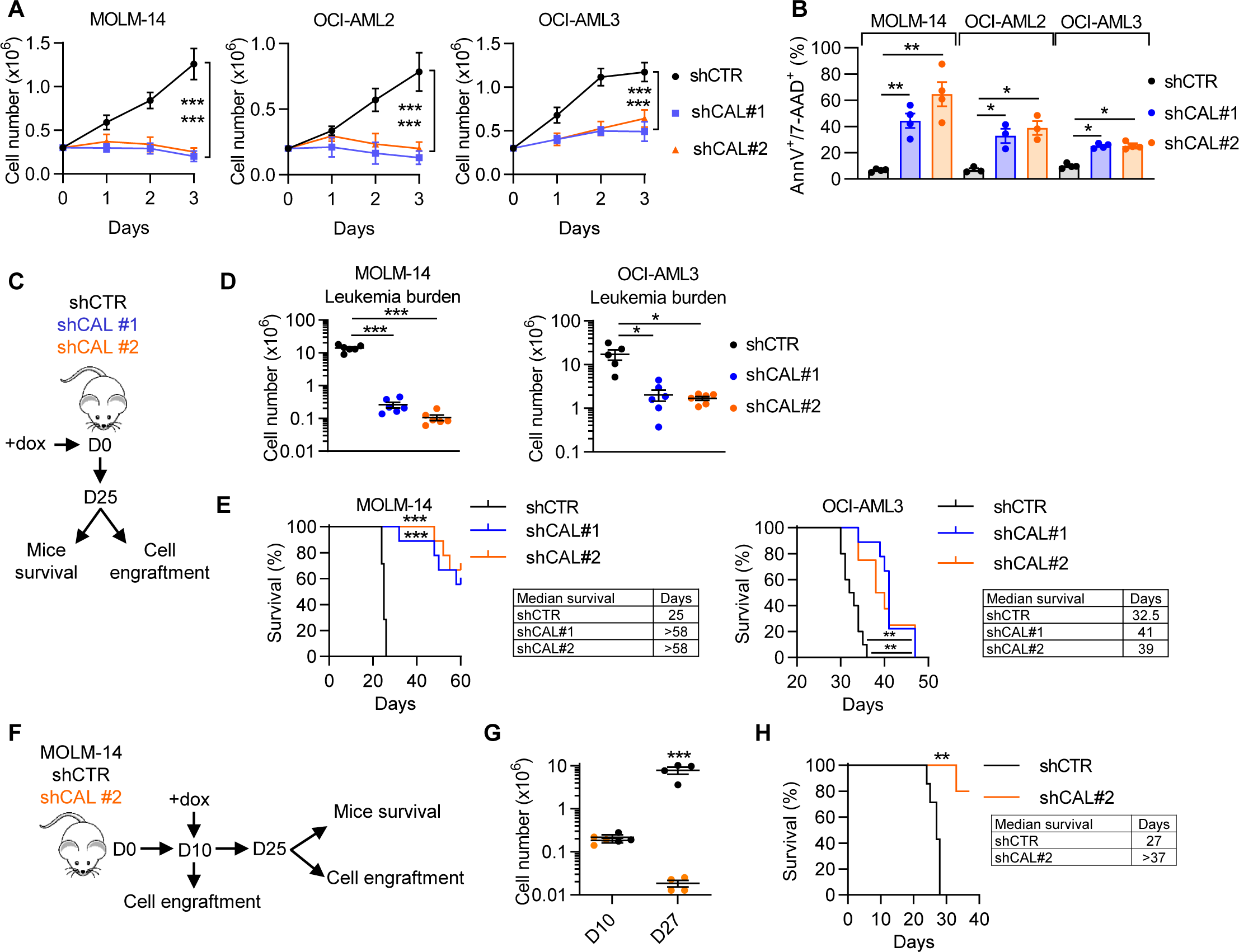
CALCRL depletion reduces AML cell growth *in vitro* and *in vivo*. (A-C) shCALCRL and control shRNA were expressed in MOLM-14, OCI-AML2 or OCI-AML3 cell lines. (A) Graph shows cell number of MOLM-14, OCI-AML2 or OCI-AML3. Three days after transduction, cells were plated at 0.3M cells per ml (D0) and cell proliferation was followed using trypan blue exclusion. (B) Graph shows the percentage of Annexin-V^+^ or 7-AAD^+^ cells 4 days after cell transduction. (C) Investigation of the role of CALCRL on leukemic cell growth *in vivo*. 2.10^6^ MOLM-14 or OCI-AML3 cells expressing doxycycline-inducible shCTR, shCAL#1 or shCAL#2 were injected into the tail vein of NSG mice. The day of injection, shRNA expression was induced adding 0.2 mg/ml doxycycline in drinking water supplemented with 1% sucrose until the end of experiment. After 25 days, a part of mice were sacrificed to measure cell engraftment (leukemia burden = blasts number into bone marrow + spleen; n=5-6 mice per group) and for another part of mice the survival parameter was followed (n=7-9 mice per group). (D) Leukemia burden measured using mCD45.1^-^/hCD45^+^/hCD33^+^/AnnV^-^ markers. (E) Mice survival monitoring. (F) Investigation of the role of CALCRL on leukemic cell growth *in vivo* in a context of engrafted cells. 2.10^6^ MOLM-14 cells expressing shCTR or shCAL#2 were injected into blood of mice. After confirmation of bone marrow cell engraftment on day 10, shRNA expression was activated by the adding of doxycycline. Then, mice survival and cell engraftment parameters were followed as previously described. (G) Measurement of Leukemia burden. (H) Mice survival monitoring. *p<0.05; **p<0.01; ***p<0.001; ns, not significant.

### CALCRL is required for Leukemic Stem Cell maintenance

As *CALCRL* expression is linked to an immature phenotype and CALCRL depletion impaired AML cell growth, we next aimed to address the role of CALCRL in LSC biology. First, we analyzed previously published single cell RNA-sequencing data (van Galen et al., 2019) and observed that *CALCRL* is preferentially expressed in HSC-like and progenitor-like cells (Prog-like cells) compared to more committed cells in 11 AML patients (Figure 3A-B-C). Moreover, while HSC-like and Prog-like cells represent only 34.3% of the total of leukemic cells found in these patients, they accounted for more than 80% of CALCRL^positive^ cells (Figure 3D). Gene Set Enrichment Analysis (GSEA) confirmed that several LSC-associated gene signatures (Eppert et al., 2011; Gentles, 2010; Ng et al., 2016) (Table S4) are significantly enriched in AML patients (the Cancer Genome Atlas, AML cohort, 2013) exhibiting the highest *CALCRL* expression compared to AML patients with the lowest *CALCRL* expression (Figure 3E). To functionally investigate the role of CALCRL in LSC biology, we performed *ex vivo* assays knocking down CALCRL in 3 primary AML samples followed by Limiting Dilution Assay (LDA). We demonstrated that in all tested samples CALCRL inhibition significantly decreased the frequency of LSCs (Figure 3F-G; see Figure S4A for gating strategy), demonstrating the requirement of CALCRL in preserving the function of LSCs.

**Figure 3.**
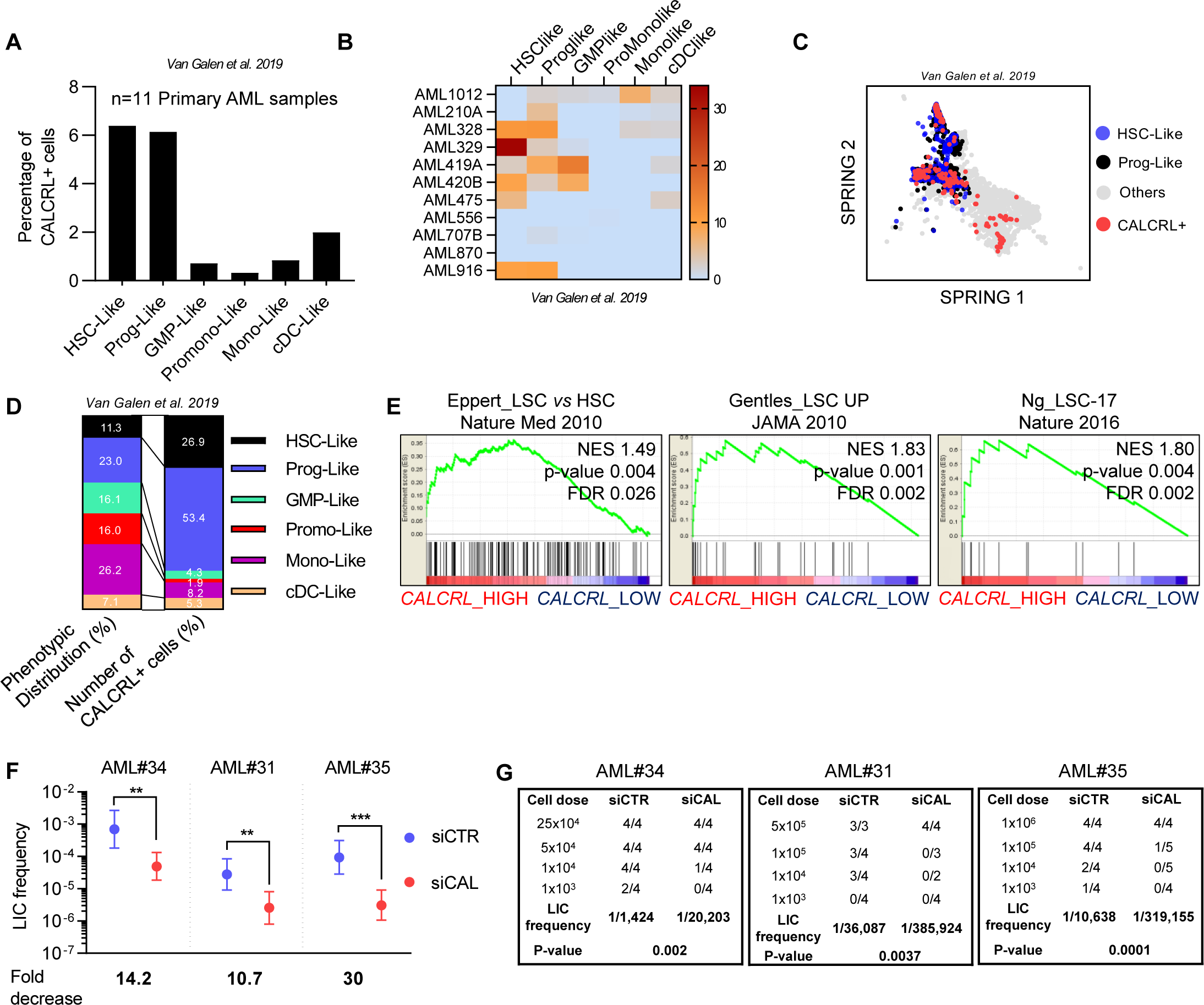
CALCRL is required for leukemic stem cell maintenance. (A) Barplot shows the percentage of CALCRL^positive^ cells in (n=11 primary AML samples) in cells classified as HSC-like, progenitor-like, GMP-like, Promonocyte-like, Monocyte-like, or cDC-like malignant cells (Van Galen et al. 2019). (B) Heatmap shows the percentage of CALCRL^positive^ cells in each patient individually. (C) SPRING visualization of single-cell transcriptomes. Points are color-coded by indicated cell-type annotations. (D) Phenotypic distribution and number of CALCRL^positive^ cells. (E) GSEA of stem cell signatures functionally identified by Eppert et al. or phenotypically defined by Gentles et al. or Ng et al. was performed using transcriptomes of cells expressing low (blue) vs high (red) levels of CALCRL gene (TCGA, AML cohort). (F) Primary AML samples or cells from primary mice were collected and treated *ex vivo* with siCTR or siCALCRL and transplanted in limiting doses intro primary or secondary recipients. Human marking of >0.1% was considered positive for AML engraftment except for AML#31 for which the cut-off was 0.5% because the sample was hCD33-. (G) Engraftment results. *p<0.05; **p<0.01; ***p<0.001; ns, not significant.

### Depletion of CALCRL alters cell cycle and DNA repair pathways in AML

To examine regulatory pathways downstream of CALCRL, we generated and performed comparative transcriptomic and functional assays on shCTR *vs* shCALCRL MOLM-14 cells. CALCRL knockdown was associated with a significant decrease in the expression of 623 genes and an increase of 278 (FDR<0.05) (Figure 4A; Table S6). Data mining and western blotting analyses showed significant depletion in genes involved in cell cycle and DNA integrity pathways (Figures 4B-C; Table S3) and a reduction in protein level of RAD51, CHEK1 and BCL2 in shCALCRL AML cells (Figure 4D). This was associated with an accumulation of cells in the G_1_ phase (Figure 4E-F). Interestingly, enrichment analysis showed that depletion of CALCRL affects the gene signatures of several key transcription factors such as E2F1, P53 or FOXM1 described as critical cell cycle regulators (Figure 4G). We focused on the E2F1 transcription factor, whose importance in the biology of leukemic stem/progenitors cells has recently been shown (Pelicano et al., 2018). We first confirmed that CALCRL depletion was closely associated with a significant decrease in the activity of E2F1 (Figure 4H). As E2F1 activity is mainly regulated by CDK-cyclin complexes, we assessed the protein expression of these actors after CALCRL downregulation. We observed a decrease in both the phosphorylation of Rb and the expression of cyclins A, B1, D1, E, reflecting the cell cycle arrest in cells depleted for CALCRL (Figure 4I).

**Figure 4.**
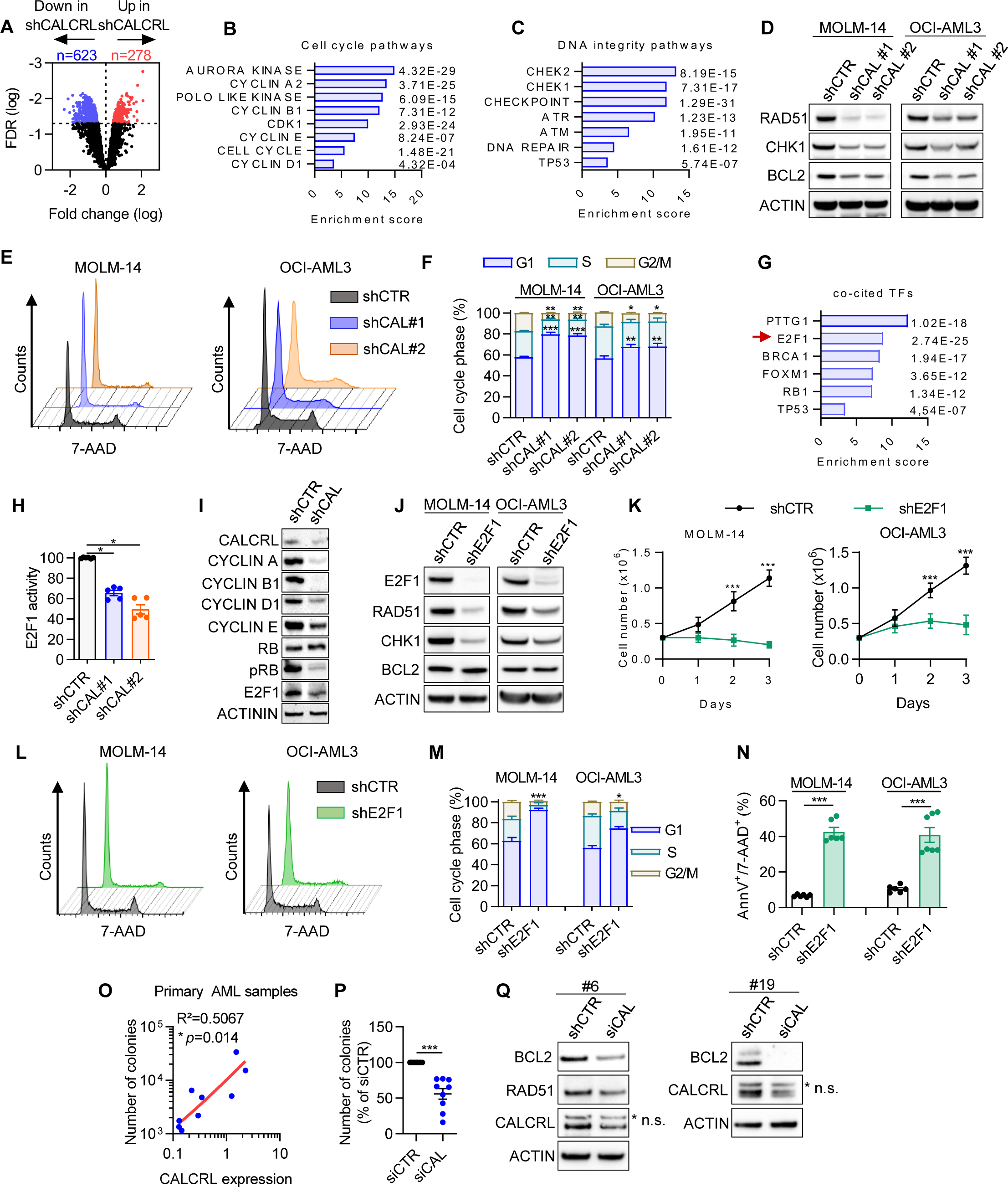
CALCRL controls cell cycle and DNA repair pathways through E2F1. (A) Volcano plot of most differentially expressed (278 upregulated and 623 downregulated) genes identified in transcriptomes of MOLM-14 shCAL#1 and #2 *vs* shCTR. The FDR values based on −log10 were plotted against the log2 ratio of gene expression level for all genes. (B-C) Genomatix analysis of cell cycle (B) or DNA integrity (C) pathways negatively affected by CALCRL depletion. (D) Western blot results showing expression of RAD51, CHK1, BCL2 and β-ACTIN proteins in MOLM-14 and OCI-AML3 four days after transduction with indicated shRNA. (E) Representative graphs of cell cycle. (F) Cell cycle distribution. (G) Genomatix analysis of co-cited transcription factors negatively affected by CALCRL depletion. (H) E2F1 activity is followed by flow cytometry through the level of mKate2 fluorescence intensity. (I) Western blot results showing expression of Cyclin A, B1, D1, E, RB and p-RB. (J) Western blot results showing expression of E2F1, RAD51, CHK1, BCL2 and -ACTIN proteins in MOLM-14 and OCI-AML3 four days after β transduction with indicated shRNA. (K) Graph shows cell proliferation of MOLM-14 or OCI-AML3. Three days after transduction, cells were plated at 0.3M cells per ml (D0) and cell proliferation was followed using trypan blue exclusion. (L) Representative graphs of cell cycle. (M) Cell cycle distribution. (N) Graph shows the percentage of Annexin-V+ or 7-AAD+ cells four days after cell transduction. (O) Correlation between clonogenic capacities of primary AML samples (n=11) and CALCRL protein expression assessed by western-blot analysis (Figure S1M). Linear regression was performed to determine R^2^ and p-value. (P) Colonies in methylcellulose were counted one week after transfection of primary AML samples with siCTR or siCAL. (Q) Western blot results showing expression of RAD51, BCL2, CALCRL and β-ACTIN proteins in primary AML samples seven days after transfection with indicated siRNA. *p<0.05; **p<0.01; ***p<0.001; ns, not significant.

Then, we demonstrated that the knockdown of E2F1 affected protein expression of RAD51, CHK1 but not BCL2 (Figure 4J), inhibited cell proliferation (Figure 4K), cell cycle progression (Figure 4L-M) and induced cell death in both MOLM-14 and OCI-AML3 (Figure 4N). We further investigated whether CALCRL might regulate the proliferation of primary AML cells. Interestingly, CALCRL protein level positively correlated with clonogenic capacities in methylcellulose (Figure 4O). Moreover, the depletion of CALCRL in primary samples decreased the number of colonies (Figure 4P), and BCL2 and RAD51 protein levels (Figure 4Q). All these results suggest that CALCRL has a role in the proliferation of AML blasts and controls critical pathways involved in DNA repair processes.

### CALCRL downregulation sensitizes leukemic cells to chemotherapeutic drugs

Based on putative targets of CALCRL such as BCL2, CHK1 or FOXM1 (David et al., 2016; Khan et al., 2017; Konopleva et al., 2016), we hypothesized that CALCRL was involved in chemoresistance. Accordingly, CALCRL depletion sensitized MOLM-14 and OCI-AML3 cells to AraC and idarubicin as attested by the reduction of cell viability (Figure 5A) and the induction of cell death (increased Annexin-V staining, Figure 5B; and increased cleavage of apoptotic proteins Caspase-3 and PARP, Figure S5A). Furthermore, depletion of ADM or E2F1 also sensitized AML cells to these genotoxic agents (Figures S5B-C), demonstrating that the ADM-CALCRL-E2F1 axis was involved in chemoresistance *in vitro*. Importantly, siRNA-mediated depletion of CALCRL in seven primary AML samples combined with AraC significantly reduced the clonogenic growth of leukemic cells compared with siCTR+AraC and siCALCRL conditions (Figure 5C). As we showed that CALCRL depletion affected DNA integrity pathways, we functionally investigated the role of CALCRL on the DNA repair by performing Comet assays. We observed that both CALCRL and ADM shRNAs increased the length of comet tails and alkaline comet tail moments compared to control cells and that AraC significantly and further increased the alkaline comet tail moment in these cells (Figures 5D-E). These results indicate that DNA repair pathways were affected when the ADM-CALCRL axis was impaired in AML.

**Figure 5.**
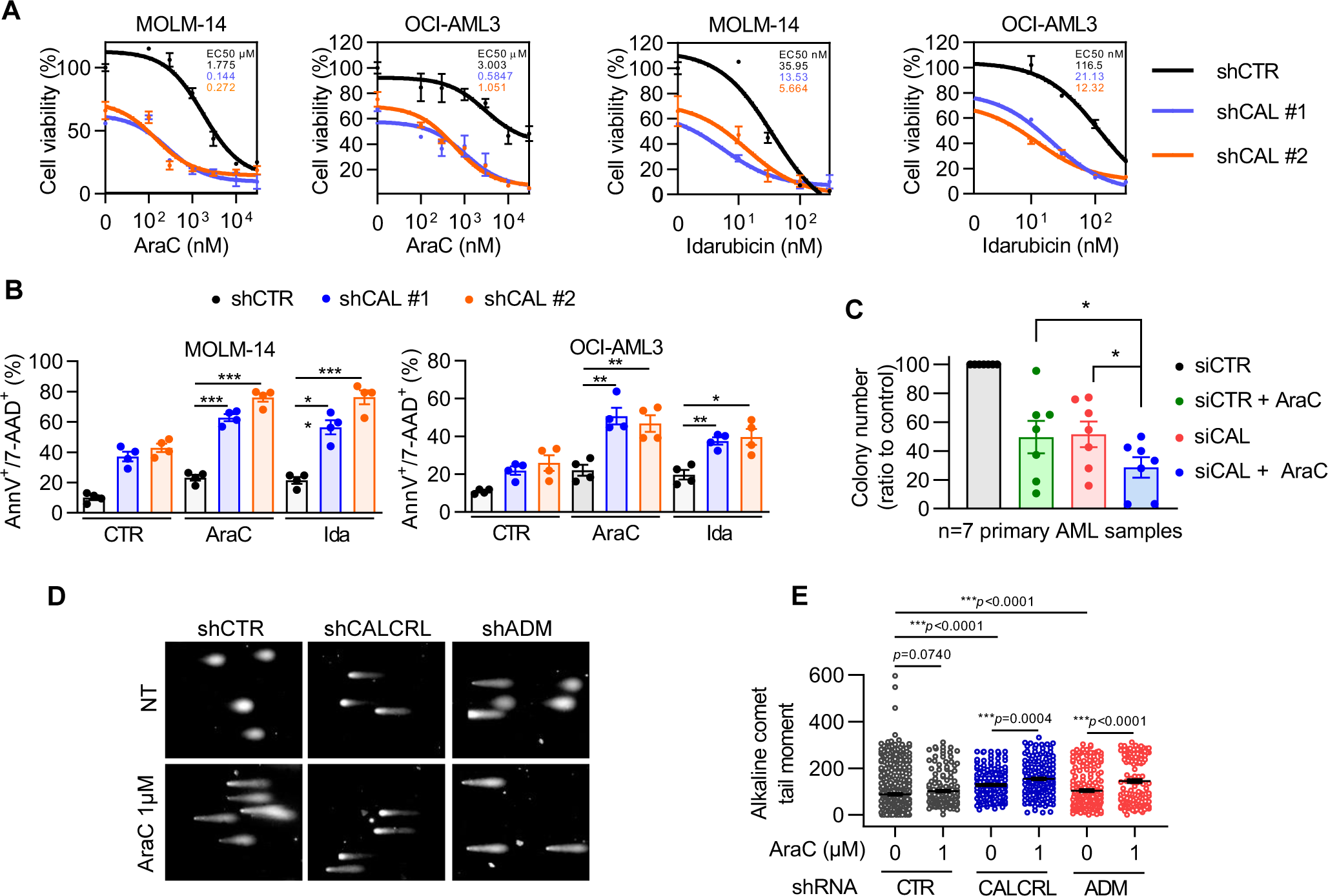
Depletion of CALCRL sensitizes AML cells to chemotherapy through induction of DNA damages. (A) Four days after transduction, MOLM-14 or OCI-AML3 cells were treated with AraC or idarubicin for 48h. Then cell viability was assessed by MTS assay and values were normalized to untreated condition. Curve fit to calculate IC50 was determined by log (inhibitor) vs. response (three parameters). (B) Graph shows the percentage of AnnV+ or 7-AAD+ cells. Four days after transduction, cells were treated with chemotherapeutic agents for 48h before flow cytometry analysis. (C) Colonies in methylcellulose were counted one week after transfection of primary AML samples with siCTR or siCAL +/-5 nM AraC (n=7 primary AML samples). (D) Detection of double strand breaks by alkaline Comet assays in MOLM-14 cells untreated or treated 24h with Ara-C (1µM). Representative pictures of nuclei in the same experiment. (E) Quantification of Alkalin Comet tail moments for one representative experiment (135–413 nuclei were analyzed for each treatment) out of four.

### Downregulation of CALCRL sensitizes leukemic cells to chemotherapy *in vivo*

We xenografted NSG mice with AML cell lines transduced with inducible shCALCRL demonstrating the same chemosensitization profile than constitutive shRNAs (Figure S5D-E). After full engraftment, CALCRL was depleted by doxycycline and mice were treated with 30 mg/kg/day AraC for 5 days (Figure 6A). While AraC alone had no effect on AML propagation, AraC in combination with shCALCRL significantly reduced the total number of blasts (Figure 6B), induced a higher rate of cell death (Figure 6C), and prolonged survival of mice (Figure 6D) compared to others conditions. Furthermore, MOLM-14 cells expressing shCTR and treated with vehicle or AraC were FACS-sorted and plated *in vitro* for further experiments. Interestingly, after one week of *in vitro* culture, human AML cells from AraC treated mice were more resistant to AraC (IC_50_: 1 μM for vehicle group *vs* 5.40 μM for AraC treated group) and idarubicin (IC_50_: 31.98 nM for vehicle group *vs* 111.3 nM for AraC treated group) (Figure 6E). Next, we observed that AML cells treated with AraC *in vivo* had higher protein expression levels of CALCRL, and a slight increase in RAD51 and BCL2, whereas CHK1 was similar to untreated cells (Figure 6F). To evaluate the role of CALCRL in this chemoresistance pathway *in vivo*, we depleted CALCRL in these cells. Knockdown of CALCRL by two different shRNAs sensitized cells to AraC and idarubicin compared to shCTR in cells treated with vehicle (Figure 6G) or AraC alone (Figure 6H). Remarkably, the IC_50_ of AraC and idarucibin in AraC-treated cells *in vivo* and transduced with shCALCRL was also decreased, demonstrating that CALCRL participated to chemoresistance pathways in AML. We further aimed to determine what ligand might be involved into this chemoresistance pathway. First, we depleted RAMP1, RAMP2 or RAMP3 using shRNAs (Figure S6A) and observed that only RAMP2 was necessary to sustain the growth of MOLM-14 cells (Figure S6B). Consistently with this result, RAMP2 downregulation induced cell death and sensitized AML cell to AraC and Ida (Figure S6C). We also showed that only exogenous ADM1-52 and ADM13-52 but not CGRP were able to decrease AraC-induced cell death in MOLM-14 cell line, highlighting that the CALCRL-driven chemoresistance is mainly due to the ADM ligand (Figure S6D). We used a PDX model treated with a CGRP inhibitor, olcegepant, in association with AraC to confirm these results (Figure S7A). We showed that CGRP inhibition did not affect leukemic burden and did not sensitize to AraC *in vivo* (Figure S7B). Interestingly, AraC upregulated CALCRL (Figure S7C-D-E), confirming our *in vitro* observations *in vivo* in PDX model. As previously described (Gluexam et al., 2019) and as a positive control of the *in vivo* activity of olcegepant, we showed that this drug efficiently decreased the percentage of murine CMP but not LSK and GMP subpopulations (Figure S7F-G-H). Altogether, these experiments indicate that ADM was the main ligand of CALCRL in AML.

**Figure 6:**
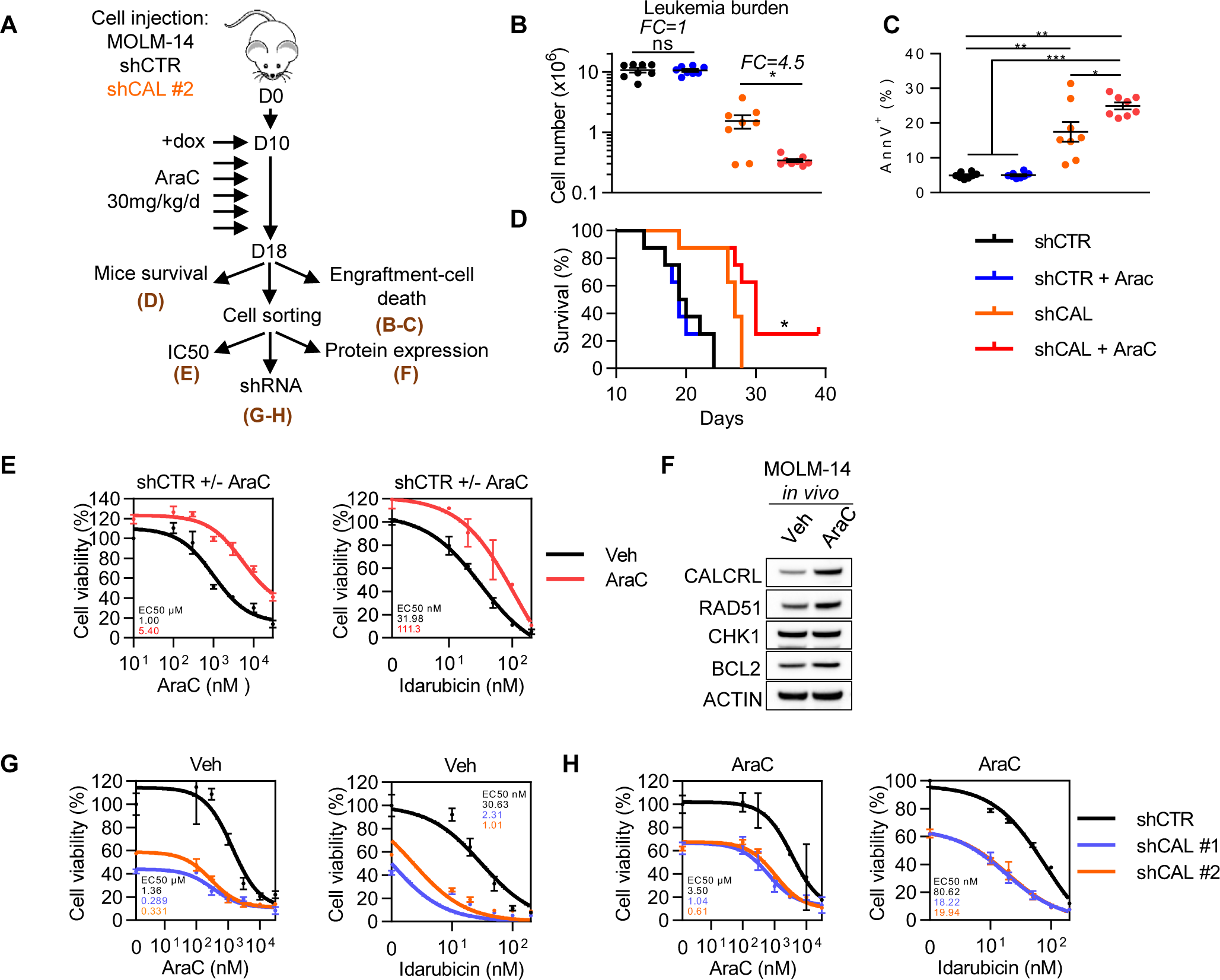
Depletion of CALCRL sensitizes AML cells to chemotherapy *in vivo*. (A) Experimental plan for assessing the consequence of CALCRL depletion on chemotherapy response *in vivo*. 2.10^6^ MOLM-14 expressing indicated inducible shRNAs were injected into the tail vein of NSG mice. Ten days later, when disease was established, mice were treated with 30mg/kg/d AraC for five days. On day 18, a group of mice was kept to follow survival (D). The other group was sacrificed to assess human cell engraftment (B) and cell death (C). Human cells were also sorted by flow cytometry using mCD45.1^-^/hCD45^+^/hCD33^+^/AnnV^-^ markers and replated in culture dishes to assess protein expression (F), sensitivity to chemotherapeutic drugs (E) and to perform CALCRL depletion by shRNA (G-H). (B) Total leukemia burden measured using mCD45.1^-^/hCD45^+^/hCD33^+^/AnnV^-^ markers. (C) Graph shows the percentage of AnnV^+^ cells. (D) Mice survival monitoring. (E) After replating, cells were treated with AraC or idarubicin *ex vivo* for 48h. Cell viability was assessed by MTS assay and values were normalized to untreated condition (n=1 in quadriplate). (F) Western-Blotting for CALCRL, RAD51, CHK1, BCL2 and β-ACTIN. Cellular extracts were collected 2 days after mice sacrifice. (G-H) Human cells from vehicle (G) or AraC (H) treated mice were transduced with indicated shRNA. After four days cells were treated with AraC or idarubicin for 48h and cell viability was assessed by MTS assay (n=1 in quadriplate). Curve fit to calculate IC50 was determined by log (inhibitor) vs. response (three parameters) test. *p<0.05; **p<0.01; ***p<0.001; ns, not significant.

Because mitochondrial metabolism has emerged as a critical regulator of cell proliferation and survival in basal and chemotherapy-treated conditions in AML (Li et al 2019; Molina et al., 2018; Scotland et al., 2013; Sriskanthadevan et al., 2015; Farge et al., 2017; Laganidou et al 2013), we analyzed the impact of CALCRL depletion on mitochondrial function. GSEA showed a significant depletion in the gene signature associated with mitochondrial oxidative metabolism in the shCALCRL MOLM-14 cells (Figure S8A; Table S3). Mitochondrial oxygen consumption measurements revealed a modest but significant reduction in basal OCR, whereas maximal respiration was conserved, indicating that mitochondria remain functional (Figure S8B). We also consistently observed a significant decrease in mitochondrial ATP production by shCALCRL (Figure S8C). We and other groups have previously shown that chemoresistant cells have elevated oxidative metabolism and that targeting mitochondria in combination with conventional chemotherapy may represent an innovative therapeutic approach in AML (Lagadinou et al 2013; Farge et al., 2017; Kuntz et al., 2017). Since depletion of CALCRL modestly decreased OCR and more greatly decreased mitochondrial ATP in AML cells (Figure S8B-C), we assessed cellular energetic status associated with AraC. Knockdown of CALCRL significantly abrogated the AraC-induced increase of basal respiration and maximal respiration (Figure S8D). Moreover, we observed a decrease in mitochondrial ATP production in response to AraC upon CALCRL silencing (Figure S8E) whereas glycolysis (e.g. ECAR) was not affected (Figure S8F).

As it has been reported that BCL2 controlled oxidative status in AML cells (Lagadinou et al., 2013), we investigated its role downstream of CALCRL. We showed that upon AraC treatment, the overexpression of BCL2 in MOLM-14 cells (Figure S8G) is sufficient to rescue maximal respiration but not basal respiration. This suggested a role of the CALCRL-BCL2 axis in maintaining some aspects of mitochondrial function in response to AraC. This was not related to energy production, as neither mitochondrial ATP production nor ECAR were affected (Figure S8I-J). Finally, BCL2 rescue almost entirely inhibited basal apoptosis induced by the depletion of CALCRL and by the combination with AraC or idarubicin (Figure S8K).

Overall, these results suggest that CALCRL mediates the chemoresistance of AML cells.

### Depletion of CALCRL in residual disease after AraC treatment impedes LSC function

To address the role of CALCRL in response to chemotherapy in primary AML samples, we used a clinically relevant PDX model of AraC treatment in AML (Farge et al., 2017). After engraftment of primary AML cells, NSG mice were treated for 5 days with AraC and sacrificed at day 8 to study the minimal residual disease (MRD; Figure 7A). We analyzed 10 different PDX models and stratified them according to their response to AraC as low (fold change, FC AraC-to-Vehicle <10) or high (FC>10) responders (Figure 7B). The percentage of cells positive for CALCRL in the AML bulk was approximately doubled in the low responder group compared to the high responder group (3.6% *vs* 7.8%; Figure 7C). We also observed an inverse correlation between the percentage of positive cells and the degree of tumor reduction (p=0.0434; Figure 7D). Moreover, after AraC treatment a significant increase in the percentage of blasts positive for CALCRL was observed (5.6% *vs.* 23%; Figure 7E) in all CD34/CD38 subpopulations (Figure 7F) from MRD. We next investigated the effects of AraC on ADM secretion. To this end, we evaluated the protein level of ADM in bone marrow supernatants of mice treated with PBS or AraC. Chemotherapy reduced both percentage of human cells (Figure S9A) and levels of secreted ADM (Figure S9B) in the bone marrow of mice. The correlation between leukemia burden and secretion of ADM reinforced the hypothesis of an autocrine secretion of ADM by leukemic blasts.

**Figure 7.**
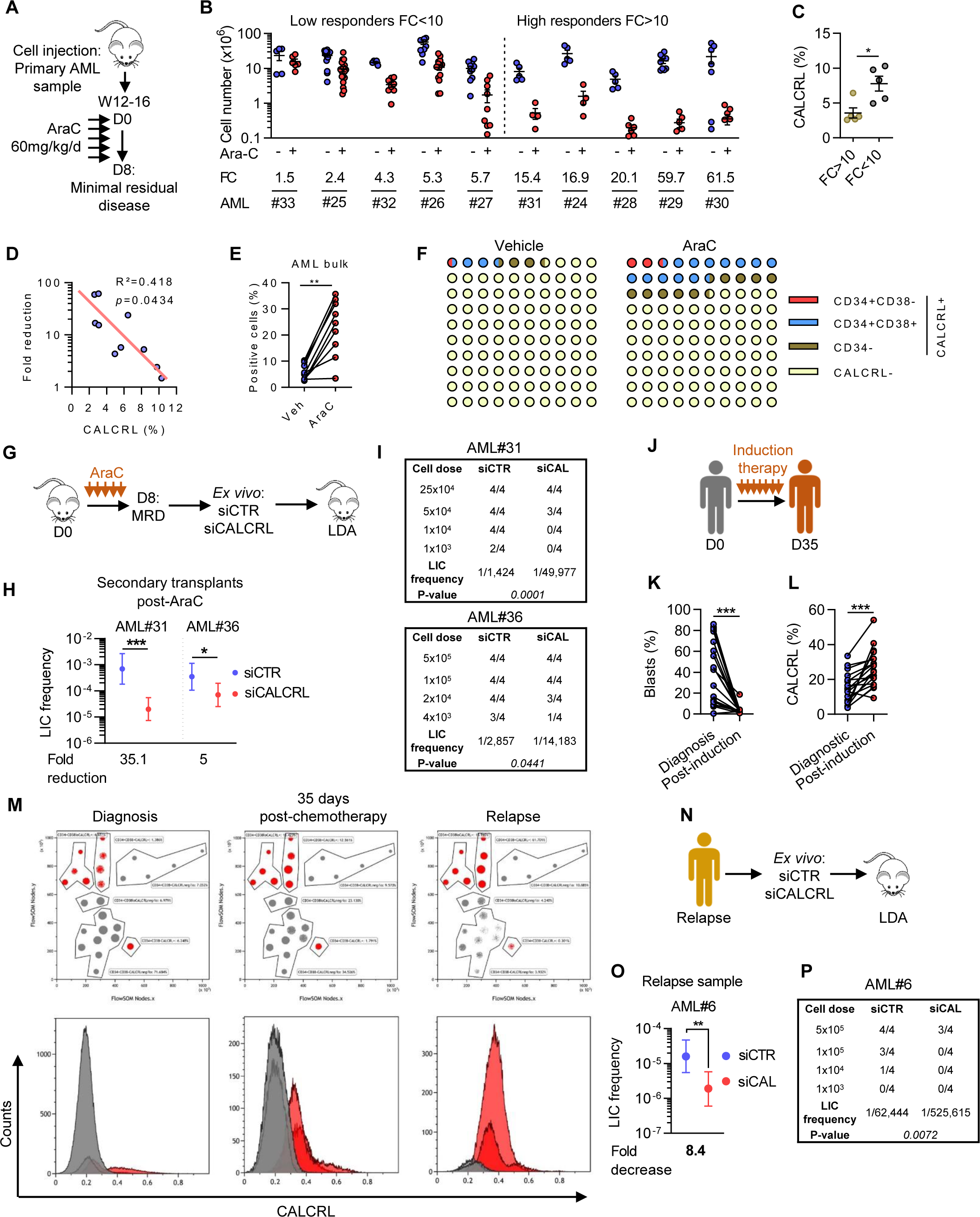
CALCRL predicts response to chemotherapy in PDX models and is required for the maintenance of R-DTCs. (A) Schematic diagram of the chemotherapy regimen and schedule used to treat NSG-based PDX models with AraC. Peripheral blood engraftment was assessed between 8 and 18 weeks, and mice were assigned to experimental groups of 4 to 10 mice with similar average engraftment per group. Mice were treated with vehicle (PBS) or 60 mg/kg/day AraC given daily *via* intraperitoneal injection for 5 days. Mice were sacrificed post treatment at day 8 to characterize viable residual AML cells. (B) Total number of human viable AML cells expressing hCD45, hCD33, and/or hCD44 was analyzed and quantified using flow cytometry in AraC-treated AML-xenografted mice compared with PBS-treated AML-xenografted mice in bone marrow. Fold reduction of leukemia burden in AraC-treated mice compared with control-treated mice was calculated individually for each AML patient sample. Patients were then separated into two categories: low responders (FC>10) and high responders (FC>10). (C) Graphs shows the percentage of cells positive for CALCRL (cell surface expression was determined by flow cytometry analysis of vehicle-treated cells) in low *vs* high responder groups. (D) Correlation between the fold reduction and the percentage of CALCRL-positive cells. (E) Percentage of human CALCRL^positive^ cells in vehicle-treated *vs* AraC-treated mice was determined by flow cytometry. (F) Picture representing the proportion of CALCRL-positive cells before and after AraC treatment according to the studied cell population. (G) Role of CALCRL in the Residual-LSC population. Primary AML sample was intravenously injected into mice. Then animals were treated with vehicle (PBS) or 60 mg/kg/day AraC given daily *via* intraperitoneal injection for 5 days. Mice were sacrificed post treatment at day 8, and siRNA transfection was performed *ex vivo* on human cells from PBS and AraC treated conditions. Decreasing cell concentrations were injected into the tail vein of mice. When engraftment was validated, mice were dissected and human cell engraftment assessed in the murine bone marrow using mCD45.1-/hCD45+/hCD33+/AnnV-markers. Human marking of >0.1% was considered positive for AML#36 sample and of 0.5% for AML#31 because the sample was hCD33-. (H) Graph shows the LIC frequency into bone marrow. (I) Engraftment results. (J) Monitoring of CALCRL expression at patient diagnosis or 35 days after intensive chemotherapy. (K) Percentage of blasts in patient bone marrow at diagnosis or after induction. (L) Percentage of human CALCRL^positive^ cells in patients was determined by flow cytometry. (M) Distribution of CALCRL^positive^ and CALCRL^negative^ cells at diagnosis, after chemotherapy, or at relapse. (N) Role of CALCRL in the LSC population at the time of relapse. After CALCRL depletion, primary AML sample was intravenously injected into mice and when engraftment was confirmed, we measured the percentage of human cells in the murine bone marrow. (O) Graph shows the LIC frequency into bone marrow. (P) Engraftment results. *p<0.05; **p<0.01; ***p<0.001; ns, not significant.

Next, we aimed at determining the role of CALCRL in the LSC function maintenance of the RIC population. Leukemic cells from patients collected at diagnosis were injected into NSG mice, and after engraftment and treatment with AraC, human viable AML cells constituting minimal residual disease were collected and transfected with siCTR or siCALCRL before LDA transplantations into secondary recipients (Figure 7G). A significant reduction of LSC frequency was observed in samples depleted from CALCRL compared to the controls in the two primary AML samples tested (Figure 7H-I). We next examined cell surface expression of CALCRL in patients before and after intensive chemotherapy (Figure 7J). Treatment decreased the percentage of blasts in the bone marrow (Figure 7K), accompanied with a significant enrichment in CALCRL^positive^ cells (Figure 7L). Moreover, we observed a continuous enrichment in CALCRL^positive^ blasts following chemotherapy (12.9% at diagnosis, 32.8% at day 35, 81% at relapse; Figure 7M). We further performed an *ex vivo* assay on a relapse sample followed by LDA in NSG mice (Figure 7N). We demonstrated that CALCRL depletion significantly reduced LSC frequency, highlighting a critical role for CALCRL in the maintenance of the clone present at relapse (Figure 7O-P). Recently Shlush *et al*. proposed an elegant model of relapses with two situations: in the first one called “relapse origin-primitive” (ROp), relapse originated from rare LSC clones only detectable in HSPC or after xenotransplantation. In the second model, called “relapse origin-committed” (ROc), the relapse clone arose from immunophenotypically committed leukemia cells in which bulk cells harbored a stemness transcriptional profile (Shlush et al., 2016). We analyzed this transcriptomic database and found that at the time of diagnosis, *CALCRL* expression was higher in blasts with ROc than with ROp phenotype, in accordance with the expression of *CALCRL* in cells harboring stem cell features (Figure S9C). Interestingly, CALCRL was strongly increased at relapse in ROp patients, which correlated with the emergence of a clone with stem cell properties at this stage of the disease (Figure S9D). These observations supported the hypothesis of the preexistence of a relapse-relevant LSC population, rare (ROp) or abundant (ROc), expressing high levels of *CALCRL*.

Altogether, these results strongly support the conclusion that CALCRL preserves LSC function after chemotherapy, thus representing an attractive therapeutic target to eradicate the clone at the origin of relapse.

## Discussion

LSC-selective therapies represent an unmet need in AML due to high plasticity and heterogeneity not only of the phenotype (Taussig et al., 2010; Eppert et al., 2011; Sarry et al., 2011) but also for the drug sensitivity (Farge et al., 2017; Boyd et al., 2018) of the LSC population. However, fundamental studies focusing on intrinsic properties of LSCs such as their resistance to chemotherapy are crucially needed for the development of better and more specific therapies in AML.

Our study provides key insights of LSC biology and drug resistance and identifies the ADM receptor CALCRL as a master regulator of RICs. Our work first shows that *CALCRL* gene is overexpressed in the leukemic compartment compared to normal hematopoietic tissue based on Eppert’s study that functionally characterizes LSCs. CALCRL could be specifically upregulated by LSC-related transcription factors such as HIF1α or ATF4 (Wang et al., 2011; van Galen et al., 2018). Indeed, both *ADM* and *CALCRL* possess the consensus hypoxia-response element (HRE) in the 5’-flanking region and are HIF1α-regulated genes (Nikitenko et al., 2003). Recently, it has been demonstrated that the integrated stress response and the transcription factor ATF4 is involved in AML cell proliferation and is uniquely active in HSCs and LSCs (van Galen et al., 2018; Heydt et al., 2018). Interestingly, maintenance of murine HSCs under proliferative stress but not steady-state conditions is dependent on CALCRL signaling (Suekane et al., 2019). Accordingly, CALCRL might support leukemic hematopoiesis and overcome stress induced by the high proliferation rate of AML cells. As opposed to ADM, CGRP is slightly expressed in BM from AML patient biopsies at diagnosis and its expression does not have a prognostic value in AML. Under inflammatory conditions to regulate immune responses or post-chemotherapeutic stress responses, continuous CGRP secretion is increased in normal hematopoietic cells or relapse-initiated cells as paracrine signal (Suekane et al., 2019; Gluexam et al., 2019). In this context, CGRP is one of the components of the CALCRL effects, while the ADM-CALCRL is the major driver of these RICs in autocrine dependent manner.

Our findings clearly show that targeting CALCRL expression impacts clonogenic capacities, cell cycle progression and genes related to DNA repair and genomic stability. Cancer stem cells and LSCs are predominantly quiescent thereby spared from chemotherapy. However, recent studies suggested that LSCs may also display a more active cycling phenotype (Iwasaki et al., 2015; Pei et al., 2018). C-type lectin CD93 is expressed on a subset of actively cycling, non-quiescent AML cells enriched for LSC activity (Iwasaki et al., 2015). Recently, Pei et al. showed that targeting the AMPK-FIS1 axis disrupted mitophagy and induced cell cycle arrest in AML, leading to the depletion of LSC potential in primary AML. These results are consistent with the existence of different sub-populations of LSCs that differ in proliferative state. Moreover, FIS1 depletion induces the down-regulation of several genes (e.g. CCND2, CDC25A, PLK1, CENPO, AURKB) and of the E2F1 gene signature that were also identified after CALCRL knockdown. Recently, it has been proposed that E2F1 plays a pivotal role in regulating CML stem/progenitor cells proliferation and survival status (Pellicano et al., 2018). Several signaling pathways, for instance MAPKs, CDK/cyclin or PI3K/AKT, have been described to be stimulated by ADM/CALCRL axis and may control pRB/E2F1 complex activity (Hallstrom et al., 2008; Wang et al., 1998). Other signaling mediators activated in LSCs such as c-Myc and CEBPα regulate E2F1 transcription and allow the interaction of the E2F1 protein with the E2F gene promoters to activate genes essential for DNA replication at G1/S, cell proliferation and survival in AML (Leung et al., 2008; O’Donnell et al., 2005; Rishi et al., 2014). Therefore, our analysis of cellular signaling downstream of CALCRL uncovers new pathways crucial for the maintenance and the chemoresistance of LSCs.

The characterization of RICs, which are detected at low threshold at diagnosis and strongly contributes to chemotherapy resistance, is necessary to develop new therapies with the aim of reducing AML relapses. Boyd and colleagues have proposed the existence of a transient state of Leukemic Regenerating Cells (LRC) during the immediate and acute response to AraC that is responsible for disease regrowth foregoing the recovery of LSC pool (Boyd et al., 2018). In this attractive model and in the dynamic period of post-chemotherapy MRD, CALCRL-positive AML cells belong to this LRC subpopulation and CALCRL is essential for the preservation of the LSC potential of primary chemoresistant AML cells. It would be interesting to determine whether chemotherapy rather primarily spares CALCRL positive cells and/or induces an adaptive response that increases the expression of CALCRL.

In summary, our data identify CALCRL as a new AML stem cell actor. CALCRL is involved in chemoresistance mechanisms, and its depletion sensitizes AML cells to chemotherapy *in vitro* and *in vivo*. Finally, our results pinpoint CALCRL as a new and promising candidate therapeutic target for eradicating the LSC subpopulation that initiated relapse in AML.

## Methods

### Human studies

Primary AML patient specimens are from Toulouse University Hospital (TUH), Toulouse, France]. Frozen samples were obtained from patients diagnosed with AML at TUH after signed informed consent in accordance with the Declaration of Helsinki, and stored at the HIMIP collection (BB-0033-00060). According to the French law, HIMIP biobank collection has been declared to the Ministry of Higher Education and Research (DC 2008-307, collection 1) and obtained a transfer agreement (AC 2008-129) after approbation by the Comité de Protection des Personnes Sud-Ouest et Outremer II (ethical committee). Clinical and biological annotations of the samples have been declared to the CNIL (Comité National Informatique et Libertés ie Data processing and Liberties National Committee). See Table S5 for age, sex, cytogenetics and mutation information on human specimens used in the current study.

### *In vivo* animal studies

NSG (NOD.Cg-Prkdcscid Il2rgtm1WjI/SzJ) mice (Charles River Laboratories) were used for transplantation of AML cell lines or primary AML samples. Male or Female mice ranging in age from 6 to 9 weeks were started on experiment and before cell injection or drug treatments, mice were randomly assigned to experimental groups. Mice were housed in sterile conditions using HEPA-filtered micro-isolators and fed with irradiated food and sterile water in the Animal core facility of the Cancer Research Center of Toulouse (France). All animals were used in accordance with a protocol reviewed and approved by the Institutional Animal Care and Use Committee of Région Midi-Pyrénées (France).

### Cell lines, primary cultures, Culture Conditions

For primary human AML cells, peripheral blood or bone marrow samples were frozen in FCS with 10% DMSO and stored in liquid nitrogen. The percentage of blasts was determined by flow cytometry and morphologic characteristics before purification. Cells were thawed in 37°C water bath, washed in thawing media composed of IMDM, 20% FBS. Then cells were maintained in IMDM, 20% FBS and 1% Pen/Strep (GIBCO) for all experiments.

Human AML cell lines were maintained in RPMI-media (Gibco) supplemented with 10% FBS (Invitrogen) in the presence of 100 U/mL of penicillin and 100 μg/mL of streptomycin, and were incubated at 37°C with 5% CO2. The cultured cells were split every 2 to 3 days and maintained in an exponential growth phase. All AML cell lines were purchased at DSMZ or ATCC, and their liquid nitrogen stock were renewed every 2 years. These cell lines have been routinely tested for Mycoplasma contamination in the laboratory. The U937 cells were obtained from the DSMZ in February 2012 and from the ATCC in January 2014. MV4-11 and HL-60 cells were obtained from the DSMZ in February 2012 and 2016. KG1 cells were obtained from the DSMZ in February 2012 and from the ATCC in March 2013. KG1a cells were obtained from the DSMZ in February 2016. MOLM14 was obtained from Pr. Martin Carroll (University of Pennsylvania, Philadelphia, PA) in 2011.

### Mouse Xenograft Model

NSG mice were produced at the Genotoul Anexplo platform at Toulouse (France) using breeders obtained from Charles River Laboratories. Transplanted mice were treated with antibiotic (Baytril) for the duration of the experiment. For experiments assessing the response to chemotherapy in PDX models, mice (6–9 weeks old) were sublethally treated with busulfan (30 mg/kg) 24 hours before injection of leukemic cells. Leukemia samples were thawed in 37°C water bath, washed in IMDM 20% FBS, and suspended in Hank’s Balanced Salt Solution at a final concentration of 1–10×10^6^ cells per 200 μL for tail vein injection in NSG mice. Eight to 18 weeks after AML cell transplantation and when mice were engrafted (tested by flow cytometry on peripheral blood or bone marrow aspirates), NSG mice were treated by daily intraperitoneal injection of 60 mg/kg AraC or vehicle (PBS) for 5 days. AraC was kindly provided by the pharmacy of the TUH. Mice were sacrificed at day 8 to harvest human leukemic cells from murine bone marrow. For AML cell lines, mice were treated with busulfan 20mg/kg) 24 hours before injection of leukemic cells. Then cells were thawed and washed as previously described, suspended in HBSS at a final concentration of 2×10^6^ per 200 μL before injection into bloodstream of NSG mice. For experiments using inducible shRNAs, doxycycline (0.2 mg/ml + 1% sucrose; Sigma-Aldrich, Cat# D9891) was added to drinking water the day of cell injection or 10 days after until the end of the experiment. Mice were treated by daily intraperitoneal injection of 30 mg/kg AraC for 5 days and sacrificed at day 8. Daily monitoring of mice for symptoms of disease (ruffled coat, hunched back, weakness, and reduced mobility) determined the time of killing for injected animals with signs of distress.

### Assessment of Leukemic Engraftment

At the end of experiment, NSG mice were humanely killed in accordance with European ethics protocols. Bone marrow (mixed from tibias and femurs) and spleen were dissected and flushed in HBSS with 1% FBS. MNCs from bone marrow, and spleen were labeled with anti-hCD33, anti-mCD45.1, anti-hCD45, anti-hCD3 and/or anti-hCD44 (all from BD) antibodies to determine the fraction of viable human blasts (hCD3-hCD45+mCD45.1−hCD33+/hCD44+AnnV-cells) using flow cytometry. In some experiments, we also added anti-CALCRL, anti-CD34 and anti-CD38 to characterize AML stem cells. Monoclonal antibody recognizing extracellular domain of CALCRL was generated in the lab with the help of Biotem company (France). Then antibody was labelled with R-Phycoerythrin using Lightning-Link kit (Expedeon). All antibodies were used at concentrations between 1/50 and 1/200 depending on specificity and cell density. Acquisitions were performed on a LSRFortessa flow cytometer with DIVA software (BD Biosciences) or CytoFLEX flow cytometer with CytoExpert software (Beckman Coulter), and analyses with Flowjo. The number of AML cells/μ peripheral blood and number of AML cells in total leukemia burden (in bone marrow and spleen) were determined by using CountBright beads (Invitrogen) using described protocol.

For LDA experiments, human engraftment was considered positive if at least >0.1% of cells in the murine bone marrow were hCD45+mCD45.1−hCD33+. The cut-off was increased to >0.5% for AML#31 because the engraftment was measured only based on hCD45+mCD45.1−. Limiting dilution analysis was performed using L-calc software. List of antibodies used in this work: Anti-CALCRL (Jean-Emmanuel Sarry lab), Anti-mCD45.1 PERCPCY5.5 (BD, Cat# 560580), Anti-hCD45 APC (BD, Cat# 555485), Anti-hCD34 AF700 (BD, Cat# 561440), Anti-hCD38 PECY7 (BD, Cat# 335825), Anti-hCD33 PE (BD, Cat# 555450), Anti-hCD44 BV421 (BD, Cat# 562890).

### Immunochemistry analysis

Protein expression was investigated by immunoreactivity scoring on tissue microarrays containing pretherapeutic bone marrow samples from intensively treated AML patients. Studies on the tissue microarray have been approved by the institutional review board of the University of Münster. Detailed information on the AML tissue microarray cohort and CALCRL expression has been published previously.^1^ AML tissue microarrays were stained using an anti-ADM (Abcam, ab69117) antibody as described.^2^ Briefly, following deparaffinization and heat-induced epitope unmasking, 4 µm tissue sections were incubated with the primary antibodies, followed by suitable secondary and tertiary antibodies (Dako). Immunoreactions were visualized with a monoclonal APAAP-complex and a fuchsin-based substrate-chromogen system (Dako). Counterstaining was performed with Mayer’s hemalum (Merck). Two investigators who were blinded towards patient characteristics and outcome independently assessed intensity of staining (1 = no/weak, 2 = moderate, 3 = strong staining intensity) and percentage of stained blasts at each intensity level. Subsequently, H-scores were calculated as described [H-score = 1 x (percentage of blasts positive at 1) + 2 x (percentage of blasts positive at 2) + 3 x (percentage of blasts positive at 3)].^1^ There was a good inter-investigator agreement (r = .91 for ADM, p < .0001). Samples from 179 AML patients were evaluable for CALCRL and ADM. Images were taken using a Nikon Eclipse 50i microscope equipped with a Nikon DS-2Mv.

1. Angenendt et al, Leukemia 2019. (DOI: 10.1038/s41375-019-0505-x)
2. Angenendt et al, Leukemia 2018. (DOI: 10.1038/leu.2017.208)

### Western blot analysis

Proteins were resolved using 4% to 12% polyacrylamide gel electrophoresis Bis-Tris gels (Life Technology, Carlsbad, CA) and electrotransferred to nitrocellulose membranes. After blocking in Tris-buffered saline (TBS) 0.1%, Tween 20%, 5% bovine serum albumin, membranes were immunostained overnight with appropriate primary antibodies followed by incubation with secondary antibodies conjugated to HRP. Immunoreactive bands were visualized by enhanced chemiluminescence (ECL Supersignal West Pico; Thermo Fisher Scientific) with a Syngene camera. Quantification of chemiluminescent signals was done with the GeneTools software from Syngene. List of antibodies used in this work: anti-CASPASE-3 (CST, Cat#9662), Anti-ACTIN (Millipore, Cat# MAB1501), anti-CALCRL (Elabscience, Cat# ESAP13421), anti-RAD51 (Abcam, Cat# ab133534), anti-BCL2 (CST, Cat# 2872), anti-E2F1 (C-20) (Santa Cruz, Cat# sc-193), anti-CHK1 (Santa Cruz, Cat# sc-8408), anti-RAMP1 (3B9) (Santa Cruz, Cat# sc-293438), anti-RAMP2 (B-5) (Santa Cruz, Cat# sc-365240), anti-RAMP3 (G-1) (Santa Cruz, Cat# sc-365313), anti-ADM (Thermo Fisher Scientific, Cat# PA5-24927), anti-CGRP (Abcam, Cat# ab47027), anti-PARP (Thermo Fisher Scientific, Cat# 44-698G), anti-alpha/beta-Tubulin (CST, Cat# 2148).

### Cell death assay

After treatment, 5.10^5^ cells were washed with PBS and resuspended in 200 μL of Annexin-V binding buffer (BD, Cat# 556420). Two microliters of Annexin-V-FITC (BD, Cat# 556454) and 7-amino-actinomycin D (7-AAD; Sigma Aldrich) were added for 15 minutes at room temperature in the dark. All samples were analyzed using LSRFortessa or CytoFLEX flow cytometer.

### Cell cycle analysis

Cells were harvested, washed with PBS and fixed in ice-cold 70% ethanol at −20°C. Cells were then permeabilized with 1×PBS containing 0.25% Triton X-100, resuspended in 1×PBS containing 10 µg/ml propidium iodide and 1 µg/ml RNase, and incubated for 30 min at 37°C. Data were collected on a CytoFLEX flow cytometer.

### Clonogenic assay

Primary cells from AML patients were thawed and resuspended in 100 μl Nucleofector Kit V (Amaxa, Cologne, Germany). Then, cells were nucleofected according to the manufacturer’s instructions (program U-001 Amaxa, Cologne, Germany) with 200nM siRNA scrambled (ON-TARGETplus Non-targeting siRNA #2, Dharmacon) or anti-CALCRL (SMARTpool ON-TARGETplus CALCRL siRNA, Dharmacon). Cells were adjusted to 1×10^5^ cells/ml final concentration in H4230 methylcellulose medium (STEMCELL Technologies) supplemented with 10% 5637-CM as a stimulant and then plated in 35-mm petri dishes in duplicate and allowed to grow for 7 days in a humidified CO2 incubator (5% CO2, 37°C). At day 7, the leukemic colonies (more than five cells) were scored.

### Plasmid cloning, shRNA, lentiviral production and leukemic cell transduction

shRNA sequences were constructed into pLKO-TET-ON or bought cloned into pLKO vectors. Each construct (6 μg) was co-transfected using lipofectamine 2000 (20 μL) in 10cm-dish with psPax2 (4 μg, provides packaging proteins) and pMD2.G (2 μg, provides VSV-g envelope protein) plasmids into 293T cells to produce lentiviral particles. Twenty-four hours after cell transfection, medium was removed and 10ml opti-MEM+1% Pen/Strep was added. At about 72 hours post transfection, 293T culture supernatants containing lentiviral particles were harvested, filtered, aliquoted and stored in -80°C freezer for future use. At the day of transduction, cells were infected by mixing 2.10^6^ cells in 2ml of freshly thawed lentivirus and Polybrene (Sigma-Aldrich, Cat# 107689) at a final concentration of 8 ug/ml. At 3 days post infection, transduced cells were selected using 1 μg/ml puromycin. List of plasmids used in this work: pCDH-puro-Bcl2 (Cheng et al., Addgene plasmid #46971), Tet-pLKO-puro (Wiederschain et al., Addgene plasmid #21915), CALCRL MISSION shRNA (shCALCRL#1, Sigma-Aldrich, Cat# TRCN0000356798; shCALCRL#2, Sigma-Aldrich, Cat# TRCN0000356736), E2F1 MISSION shRNA (Sigma-Aldrich, Cat# TRCN0000039659). List of shRNA sequences: shCALCRL#1 FW CTTATCTCGCTTGGCATATTC, shCALCRL#2 FW TTACCTGATGGGCTGTAATTA, shE2F1 FW CGCTATGAGACCTCACTGAAT, shRAMP1 FWCCCTTCTTCCAGCCAAGAAGA, shRAMP2 FW GAGCTTCTCAACAACCATGTT, shRAMP3 FW GGACTAGGACTCCTTGCTTGA.

### IC_50_ experiments

The day before experiment, cells were adjusted to 3×10^5^ cells/ml final concentration and plated in a 96-well plate (final volume: 100 μl). To measure half-maximal inhibitory concentration (IC_50_), increased concentrations of AraC or idarubicin were added to the medium. After two days, 20 μl per well of MTS solution (Promega) was added for two hours and then absorbance was recorded at 490nm with a 96-well plate reader. The doses that decrease cell viability to 50% (IC_50_) were analyzed using nonlinear regression log (inhibitor) vs. response (three parameters) with GraphPad Prism software.

### Measurement of oxygen consumption in AML cultured cells using Seahorse Assay

All XF assays were performed using the XFp Extracellular Flux Analyser (Seahorse Bioscience, North Billerica, MA). The day before the assay, the sensor cartridge was placed into the calibration buffer medium supplied by Seahorse Biosciences to hydrate overnight. Seahorse XFp microplates wells were coated with 50 µl of Cell-Tak (Corning; Cat#354240) solution at a concentration of 22.4 µg/ml and kept at 4°C overnight. Then, Cell-Tak coated Seahorse microplates wells were rinsed with distillated water and AML cells were plated at a density of 10^5^ cells per well with XF base minimal DMEM media containing 11 mM glucose, 1 mM pyruvate and 2 mM glutamine. Then, 180 µl of XF base minimal DMEM medium was placed into each well and the microplate was centrifuged at 80 g for 5 min. After one hour incubation at 37°C in CO_2_ free-atmosphere, basal oxygen consumption rate (OCR, as a mitochondrial respiration indicator) and extracellular acidification rate (ECAR, as a glycolysis indicator) were performed using the XFp analyzer.

### Alkalin Comet assays

Alkalin Comet assays were performed with OxiSelect Comet Assay Kit and according to the manufacturer’s instructions (Cell Biolabs Inc.). Electrophoresis was performed at 4°C in alkaline condition at 20V during 45mn. Slides were visualized by using a fluorescence microscope (AxioObserver Z1; Zeiss). Comet tail moments were measured with ImageJ software (version 1.8v) with the plugin OpenComet (http://opencomet.org/). Apoptotic cells were excluded from the analysis.

### RNA microarray and bioinformatics analyses

For primary AML samples, human CD45+CD33+ were isolated using cell sorter cytometer from engrafted BM mice (for 3 primary AML specimens) treated with PBS or treated with AraC. RNA from AML cells was extracted using Trizol (Invitrogen) or RNeasy (Qiagen). For MOLM-14 AML cell line, mRNA from 2.10^6^ of cells was extracted using RNeasy (Qiagen). RNA purity was monitored with NanoDrop 1ND-1000 spectrophotometer and RNA quality was assessed through Agilent 2100 Bionalyzer with RNA 6000 Nano assay kit. No RNA degradation or contamination were detected (RIN > 9). 100 ng of total RNA were analysed on Affymetrix GeneChip© Human Gene 2.0 ST Array using the Affymetrix GeneChip© WT Plus Reagent Kit according to the manufacturer’s instructions (Manual Target Preparation for GeneChip® Whole Transcript (WT) Expression Arrays P/N 703174 Rev. 2). Arrays were washed and scanned; and the raw files generated by the scanner was transferred into R software for preprocessing (with RMA function, Oligo package), quality control (boxplot, clustering and PCA) and differential expression analysis (with eBayes function, LIMMA package). Prior to differential expression analysis, all transcript clusters without any gene association were removed. Mapping between transcript clusters and genes were done using annotation provided by Affymetrix (HuGene-2_0-st-v1.na36.hg19.transcript.csv) and the R/Bioconductor package hugene20sttranscriptcluster.db. p-values generated by the eBayes function were adjusted to control false discovery using the Benajmin and Hochberg’s procedure. [RMA] Irizarry et al., Biostatistics, 2003; [Oligo package] Carvalho and Irizarry, Bioinformatics, 2010; [LIMMA reference] Ritchie et al., Nucleic Acids Research, 2015; hugene20sttranscriptcluster.db :MacDonald JW 2017, Affymetrix hugene20 annotation data (chip hugene20sttranscriptcluster); [FDR]: Benjamini et al., Journal of the Royal Statistical Society, 1995.

### GSEA analysis

GSEA analysis was performed using GSEA version 3.0 (Broad Institute). Gene signatures used in this study were from Broad Institute database, literature, or in-house built. Following parameters were used: Number of permutations = 1000, permutation type = gene_set. Other parameters were left at default values.

### Statistical analysis

We assessed the statistical analysis of the difference between two sets of data using two-tailed (non-directional) Student’s t test with Welch’s correction. For survival analyses, we used Log-rank (Mantel-Cox) test. Analyses were performed using Graphpad Prism (V6 and V8). For Limiting Dilution Assay experiments, frequency and statistics analyses were performed using L-calc software (Stemcell technologies). A p value of less than 0.05 indicates significance. *p < 0.05; **p < 0.01; ***p < 0.001; ****p < 0.0001; ns, not significant. Detailed information of each test is in the figure legends.

All publicly accessible transcriptomic databases of AML patients used in this study:

#### GSE30377

Eppert K, Takenaka K, Lechman ER, Waldron L, Nilsson B, van Galen P, Metzeler KH, Poeppl A, Ling V, Beyene J, Canty AJ, Danska JS, Bohlander SK, Buske C, Minden MD, Golub TR, Jurisica I, Ebert BL, Dick JE. (28 August 2011) Stem cell gene expression programs influence clinical outcome in human leukemia. Nat Med, 17(9), 1086-93.

#### GSE14468

Verhaak RG, Wouters BJ, Erpelinck CA, Abbas S, Beverloo HB, Lugthart S, Löwenberg B, Delwel R, Valk PJ. (January 2009) Prediction of molecular subtypes in acute myeloid leukemia based on gene expression profiling. Haematologica, 94(1), 131-4.

#### GSE12417

Metzeler KH, Hummel M, Bloomfield CD, Spiekermann K, Braess J, Sauerland MC, Heinecke A, Radmacher M, Marcucci G, Whitman SP, Maharry K, Paschka P, Larson RA, Berdel WE, Büchner T, Wörmann B, Mansmann U, Hiddemann W, Bohlander SK, Buske C; Cancer and Leukemia Group B.; German AML Cooperative Group. (15 November 2008) An 86-probe-set geneexpression signature predicts survival in cytogenetically normal acute myeloid leukemia. Blood, 112(10), 4193-201.

#### GSE116256

Van Galen P, Hovestadt V, Wadsworth Ii MH, Hughes TK, Griffin GK, Battaglia S, Verga JA, Stephansky J, Pastika TJ, Lombardi Story J, Pinkus GS, Pozdnyakova O, Galinsky I, Stone RM, Graubert TA, Shalek AK, Aster JC, Lane AA, Bernstein BE. Single-cell RNA-seq reveals AML hierarchies relevant to disease progression and immunity. Cell. 2019 Mar 7;176(6):1265-1281.

#### TCGA

The Cancer Genome Atlas Research Network. (30 May 2013) Genomic and epigenomic landscapes of adult de novo acute myeloid leukemia. N Engl J Med, 368(22), 2059-74. Erratum in: N Engl J Med. 2013 Jul 4;369(1):98.

#### BEAT AML

Functional genomic landscape of acute myeloid leukaemia. Nature, 2018 Oct;562(7728):526-531

## Acknowledgements

We thank all members of mice core facilities (UMS006, ANEXPLO, Inserm, Toulouse) for their support and technical assistance, and Prof. Véronique De Mas and Eric Delabesse for the management of the Biobank BRCHIMIP (Biological Resources Centres-INSERM Midi-Pyrénées “Cytothèque des hémopathies malignes”) that is supported by CAPTOR (Cancer Pharmacology of Toulouse-Oncopole and Région). We thank the cellular imaging facility of U1037-CRCT. We thank the flow cytometry core facilities of U1037-CRCT and U1048-I2MC for technical assistance, and Anne-Marie Benot, Muriel Serthelon and Stéphanie Nevouet for their daily help about the administrative and financial management of our Team RESISTAML. This work was supported by grants from the French government under the program “Investissement d’avenir” CAPTOR (ANR-11-PHUC-001), the Labex TOUCAN, the Fondation Toulouse Cancer Santé, the Fondation ARC, the Ligue National de Lutte Contre le Cancer, and from the Association G.A.E.L.

## Author contributions

Conception and design: C.L., J.-E.S. Development of methodology: C.L., J.E.S., V.F., C.R., J.T., T.K., C.S. Acquisition of data (provided animals, acquired and managed AML samples, provided facilities, etc.): C.L., M.D., P.L.M., T.F., M.G., C.B., E.S., M.-L.N.-T., M.S., N.S., Q.H., N.A., L.S., T.K., L.A., J.-H.M., S.M., N.G., E.B, A.S. Analysis and interpretation of data: C.L., J.-E.S. Study supervision: J.-E.S.

## Supplementary legends

**Figure S1. Expression of CALCRL and its ligand adrenomedullin and impact on patient outcome in AML**

(A) Impact of *CALCRL* gene expression on overall survival. (B) *CALCRL* expression (BEAT AML cohort) in patients achieving a complete remission or refractory to chemotherapy. (C) *CALCRL* expression (BEAT AML cohort) in CBF, NK and Complex AML. (D) *CALCRL* gene expression in normal *versus* leukemic CD34^+^CD38^-^, CD34+CD38low, progenitors and Lin^+^ compartments according to Eppert *et al*. (E) *CALCRL* expression (TCGA and BEAT AML cohorts) according to FAB subtypes. (F) Flow cytometry plot shows CALCRL expression on CD34+ primary AML cells. (G) Percentage of cells expressing CALCRL was assessed by flow cytometry analysis of normal CD34+ and leukemic cells. (H) Expression of *CALCRL* according to patient mutational status (AML TCGA). (I) Expression of *ADM* according to patient mutational status (AML TCGA). (J) *ADM* gene expression in normal compartments CD34+CD38-, CD34+CD38low, progenitors and Lin+ compared with leukemic LSC- and LSC+ compartments according to Eppert *et al*. (K) *ADM* gene expression in normal *versus* leukemic CD34^+^CD38^-^, CD34+CD38low, progenitors and Lin^+^ compartments according to Eppert *et al*. (L) *ADM* expression at diagnosis and relapse after intensive chemotherapy according to Hackl *et al*. (M) Western Blot results showing expression of CALCRL and ADM in normal peripheral blood mononuclear cells (n=3) or in a panel of primary AML samples (n=16). (N) Quantification of western blot. (O) Correlation between CALCRL and ADM protein expression. (P) *CALCRL*, *ADM*, *RAMP1*, *RAMP2* and *RAMP3* gene expression in AML cell lines. (Q) Western Blot results showing expression of CALCRL, RAMP1, RAMP2, RAMP3, ADM and CGRP in a panel of AML cell lines. (R) Confocal analysis of the expression of CALCRL (in green) in primary AML samples and AML cell lines. (S) *ADM*, *CALCA* and *CALCB* expression (TCGA, Verhaak *et al*. and BEAT AML cohorts).

**Figure S2. Expression of CGRP and impact on patient outcome in AML.**

(A) Representative IHC micrographs of CGRP expression in pretherapeutic BM from AML patients. (B) and (C) Overall and Event-Free Survival according to CGRP H-scores. OS and EFS were regressed against CGRP H-scores using univariate or multivariate Cox model. Data were dichotomized for visualization purpose.

*p<0.05; **p<0.01; ***p<0.001; ns, not significant.

**Figure S3. Depletion of CALCRL or ADM impairs leukemic cell growth.** (A) Western blot results showing expression of ADM and β-ACTIN proteins in MOLM-14 and OCI-AML3 four days after transduction with shADM. (B) Graph shows cell number of MOLM-14 or OCI-AML3. Three days after transduction, cells were plated (D0) and cell proliferation was followed using trypan blue exclusion. (C) Graph shows the percentage of Annexin-V+ or 7-AAD+ cells 4 days after cell transduction. (D) Western blot results showing expression of CALCRL and β-ACTIN proteins in MOLM-14 and OCI-AML3 transduced with inducible shRNAs targeting CALCRL. Protein expression was assessed 7 days after the induction of shRNA with doxycycline (1 µg/ml). (E) Graph shows cell number of MOLM-14 or OCI-AML3. Doxycycline was added at D0 and cell proliferation was followed using trypan blue exclusion. (F) Graph shows the percentage of Annexin-V+ or 7-AAD+ cells 7 days after adding doxycycline into cell culture medium. (G) Flow cytometry plots showing the gating strategy used to identify human leukemic cells issued from MOLM-14 or OCI-AML3 xenografts. (H) Picture of mice bone marrow engrafted with indicated shRNA.

*p<0.05; **p<0.01; ***p<0.001; ns, not significant.

**Figure S4. Identification of human leukemic cells issued from primary AML xenografts.** (A) Flow cytometry plots showing the gating strategy used to identify human leukemic cells issued from primary AML xenografts.

**Figure S5. Depletion of CALCRL sensitizes to chemotherapeutic drugs *in vitro*** (A) Western-Blotting for PARP, CASPASE-3, CALCRL and β-ACTIN. (B-C) Graph shows the percentage of Annexin-V+ or 7-AAD+ cells. Three days after transduction with shCTR, shADM (B), or shE2F1 (C), cells were treated with AraC or idarubicin for 48h before flow cytometry analysis. (D) Five days after adding doxycycline, MOLM-14 or OCI-AML3 cells expressing inducible shRNAs were treated with AraC or idarubicin for 48h. Then cell viability was assessed by MTS assay and values were normalized to untreated condition. Curve fit to calculate IC50 was determined by log (inhibitor) vs. response (three parameters) test. (E) Graph shows the percentage of Annexin-V+ or 7-AAD+ cells. Five days after adding doxycycline, MOLM-14 or OCI-AML3 cells expressing inducible shRNAs were treated with AraC or idarubicin for 48h. Then, the percentage of cell death was monitored using Annexin-V+ and 7-AAD+ markers.

**Figure S6. RAMP2–mediated ADM signaling drives cell proliferation and resistance to chemotherapeutic agents** (A) Expression of *RAMP1*, *RAMP2* and *RAMP3* 5 days after induction of shRNA expression with doxycycline. (B) Graph shows cell number of MOLM-14 expressing shCTR, shRAMP1, shRAMP2 or shRAMP3. Doxycycline was added at D0 and cell proliferation was followed using trypan blue exclusion. (C) Graph shows the percentage of Annexin-V+ or 7-AAD+ cells. Five days after adding doxycycline, MOLM-14 or OCI-AML3 cells expressing inducible shRNAs were treated with AraC or Idarubicin for 48h. Then, the percentage of cell death was monitored using Annexin-V+ and 7-AAD+ markers. (D) Graph shows the percentage of Annexin-V+ or 7-AAD+ MOLM-14 cells treated 48h with the indicated compound (AraC 1uM).

**Figure S7: *In vivo* inhibition of CGRP signaling in combination with chemotherapy** (A) Primary cells were injected in the tail vein of NSG and after leukemic engraftment, mice were intraperitoneally treated with olcegepant (5 mg/kg/d) alone or in combination with AraC (30 mg/kg/d). At day 11, mice were sacrificed and the minimal residual disease was studied. (B) Leukemic burden. (C) CALCRL expression in human blasts (MFI). (D) CALCRL positive cells in human blasts (%). (E) CALCRL positive cells in human CD34+CD38-cells (%). (E) Impact of treatments on murine LSKs. (F) Impact of treatments on murine CMPs. (E) Impact of treatments on murine GMPs.

**Figure S8. Downregulation of CALCRL interferes with OxPHOS status of AML cells in a BCL2-dependent manner.** (A) GSEA of the Farge_HIGHOXPHOS and HALLMARK_OXIDATIVE_PHOSPHORYLATION gene signatures in transcriptomes of MOLM-14 cells expressing shCTR (red) compared with shCALCRL (blue). (B) OCR was measured by seahorse three days after lentiviral transduction of MOLM-14 or OCI-AML3 cells with a non-targeting shRNA or a shRNA against CALCRL (shCAL#2). (C) Mitochondrial ATP production was assessed using seahorse. (D-E-F) Four days after lentiviral transduction of MOLM-14 cells with shCTR or shCALCRL (shCAL#2), cells were treated with vehicle or 2 µM AraC for 24h and OCR, mitochondrial ATP production and ECAR were measured by seahorse. (G) MOLM-14 transduced with pCDH-EMPTY or pCDH-BCL2 vectors. (H-I-J) MOLM-14 pCDH-EMPTY or pCDH-BCL2 were transduced with non-targeting shRNA or shRNA targeting CALCRL (CAL#2) for four days. Then cells were treated with 2 µM AraC for 48h before OCR, mitochondrial ATP production and ECAR measurements by seahorse. (K) Cells were exposed to 1µM AraC or 50nM Idarubicin for 48h before analysis of the percentage of AnnV+ or 7-AAD+ cells by flow cytometry.

**Figure S9. CALCRL sustains LSC function of Relapse-initiated Drug Tolerant Cells** (A) Percentage of human cells in the murine bone marrow in PBS and AraC-treated mice. (B) Western-Blot and graph showing the protein expression of ADM in the bone marrow supernatant of xenografted mice treated with PBS or AraC. (C) *CALCRL* expression in ROc and ROp patients at the time of diagnosis (Shlush et al. 2017). (D) *CALCRL* expression in ROc and ROp patients at diagnosis or relapse.

